# Clinical Antibiotic Resistance Patterns Across 70 Countries

**DOI:** 10.1101/2020.12.04.411504

**Authors:** Pablo Catalan, Carlos Reding, Jessica Blair, Ivana Gudelj, Jon Iredell, Robert Beardmore

## Abstract

We sought global patterns of antibiotic resistant pathogenic bacteria within the AMR Research Initiative database, Atlas. This consists of 6.5M clinical minimal inhibitory concentrations (MICs) observed in 70 countries in 633k patients between 2004 and 2017. Stratifying MICs according to pathogens (P), antibiotics (A) and countries (C), we found that the frequency of resistance was higher in Atlas than other publicly available databases. We determined global MIC distributions and, after showing they are coherent between years, we predicted MIC changes for 43 pathogens and 827 pathogen-antibiotic (PAs) pairings that exhibit significant resistance dynamics, including MIC increases and even decreases. However, many MIC distributions are multi-modal and some PA pairs exhibit sudden changes in MIC. We therefore analysed Atlas after replacing the clinical classification of pathogens into ‘susceptible’, ‘intermediate’ and ‘resistant’ with an information-optimal, cluster-based classifier to determine subpopulations with differential resistance that we denote S and R. Accordingly, S and R clusters for different PA pairs exhibit signatures of stabilising, directional and disruptive selection because their respective MICs can have different dynamics. Finally, we discuss clinical applications of a (R, dR/dt) ‘phase plane’ whereby the MIC of R is regressed against change in MIC (dR/dt), a methodology we use to detect PA pairs at risk of developing clinical resistance.

## Introduction

Antibiotic resistance is a multiscale phenomenon. Small molecule antibiotics bind sites measurable on the nanoscale and yet those molecules, the pathogens they target and their resistance genes traverse our planet. The problem of predicting global changes in antibiotic resistance and its economic burden is, therefore, particularly difficult. Governments have attempted this, predicting future increases in resistance, disease morbidity and mortality^*1,2*^ but, possibly because of low fidelity and sparse antibiotic usage and resistance data, long-term extrapolations made from narrow ranges of clinical conditions have been criticised.^*3*^

One could, therefore, simplify the question and ask how antibiotic consumption correlates with resistance. Self-evidently, reduced consumption should reduce resistance^*2,4*^ but this is surprisingly difficult to establish. One problem may be timescales: theory predicts resistance rises faster than it decays following antibiotic withdrawal^*5*^ and, in practise, the removal of an antibiotic is known not to always lead to reductions in resistance, for instance at the scale of hospitals.^*6,7*^ However, resistance has reduced after city-scale interventions^*8*^ and following pathogen-specific interventions.^*9*^ Already amplified resistance genes can be rapidly lost from microbial genomes following drug withdrawal^*10*^ but resistant species need not rescind from microbial communities.^*11*^ At the scale of clinical patients, longitudinal pathogen isolates can exhibit volatile resistance patterns, like a 500-fold weekly change in antibiotic susceptibility that includes cross-resistance to antibiotics not used for treatment.^*12,13*^ However, such volatility may result from sampling low cell numbers or culturing clones that are not representative of the wider pathogen population.

Given such complexities, whether we can reverse resistance^*14–16*^ systematically is a difficult open problem and as a means to assess resistance changes, ideally reductions, high fidelity spatiotemporal data on the *status quo* is needed. The Wellcome Trust and Open Data Institute MIC database project, *Atlas*,^*17*^ seeks exactly this, so we compared Atlas minimal inhibitory concentration (MIC) data to other resistance databases and made predictions on which pathogen-antibiotic (PA) pairs exhibit the greatest changes in resistance.

Each Atlas datum is a vector of MICs determined for a pathogen isolated from one patient where each MIC completely inhibits the growth of a pathogen in an antibiotic susceptibility assay. Pathogens are classified as resistant and an antibiotic is not recommended for treatment if the MIC lies above a prior clinical breakpoint.^*18–20*^ The MIC is a standardised, albeit variable, even laboratory- and assay-dependent^*21*^ resistance measure and despite the inevitable noise and anomalies within Atlas, we sought signals in the data collated over a decade-long period (2004-2017). This represents 6.5M MICs for pathogens isolated from approximately 633k patients in 70 countries.

The database holds data for 284 pathogens but only those represented by more than 500 entries and with temporal data from more than two years were included in an initial sift, meaning 43 pathogens and 827 pathogen-antibiotic pairs out of 3,919 in the database. We will show many of their MIC distributions exhibit hallmarks of phenotypic evolution: multi-modal distributions with subpopulations that cluster around different MICs that shift each year. Some of these are consistent with purifying, directional and disruptive selection acting on MICs and we detail these patterns for important clinical pathogens below.

## Results

### Testing bias towards low MICs

We were first concerned Atlas data could bias towards low MICs following years in which pathogen sampling methods changed. For instance, could heightened awareness of resistance lead to increased clinical susceptibility testing that could, in turn, increase the reporting of low MICs? Could improvements in molecular identification methodologies^*22,23*^ have an analogous effect? To address this we sought PA pairs that initially exhibited increasing MIC until a significant increase in testing led to a concomitant decrease in MIC such that the overall MIC change was negative. Now, from all possible PA pairs, irrespective of data quantity, 724 report significantly decreasing mean MICs (although we show below that these may still have subpopulations with increasing clinical resistance) and 481 of these are concomitant with increased MIC testing. Of those, a changepoint analysis finds 13 pairs exhibit a significant change in MIC derivative (from increasing to decreasing; Figure S1) and these, we hypothesise, are most likely to be affected by sampling bias towards low MIC values. This suggests testing-induced low-MIC bias is present and other methods may uncover additional biases. These may be sufficiently rare so as to not affect database-wide analyses but biases must be accounted for when a rationale for MIC change is applied to any PA pair and some will be discussed below.

### Between-database consistency: Atlas, ResistanceMap and ECDC

Atlas curators use different sources, they acknowledge variability between sources and address some inconsistencies.^*17*^ We summarise basic statistics (Figure S2) which indicate increases in data quantity through time and how US data, *Staphylococcus aureus* and *Escherichia coli* data dominate. Importantly, Atlas contains raw, anonymised patient MICs and metadata whereas other programmes typically report proportions of resistance longitudinally, like the Public Health England ESPAUR report.^24^ There, the use of an essentially binary filter based on clinical breakpoints limits the analyses that can be done and, due to the way data is published, we could not compare UK data in Atlas with ESPAUR. Now, applying the clinical breakpoint classifier to Atlas and comparing against the most recent European Centre for Disease Prevention and Control (ECDC)^*25*^ and ResistanceMap^*26*^ databases we find Atlas typically reports higher frequencies of resistance (Figures 1A, S3). Moreover, between-database point differences are as large as 60% for some PA pairs (Figure 1A) where within-country correlations are high for France and Portugal but low for Denmark, Netherlands and others (Figure S3). This may be explained by smaller databases having larger between-database differences (Figure S4, *p* < 0.002).

**Fig. 1:**
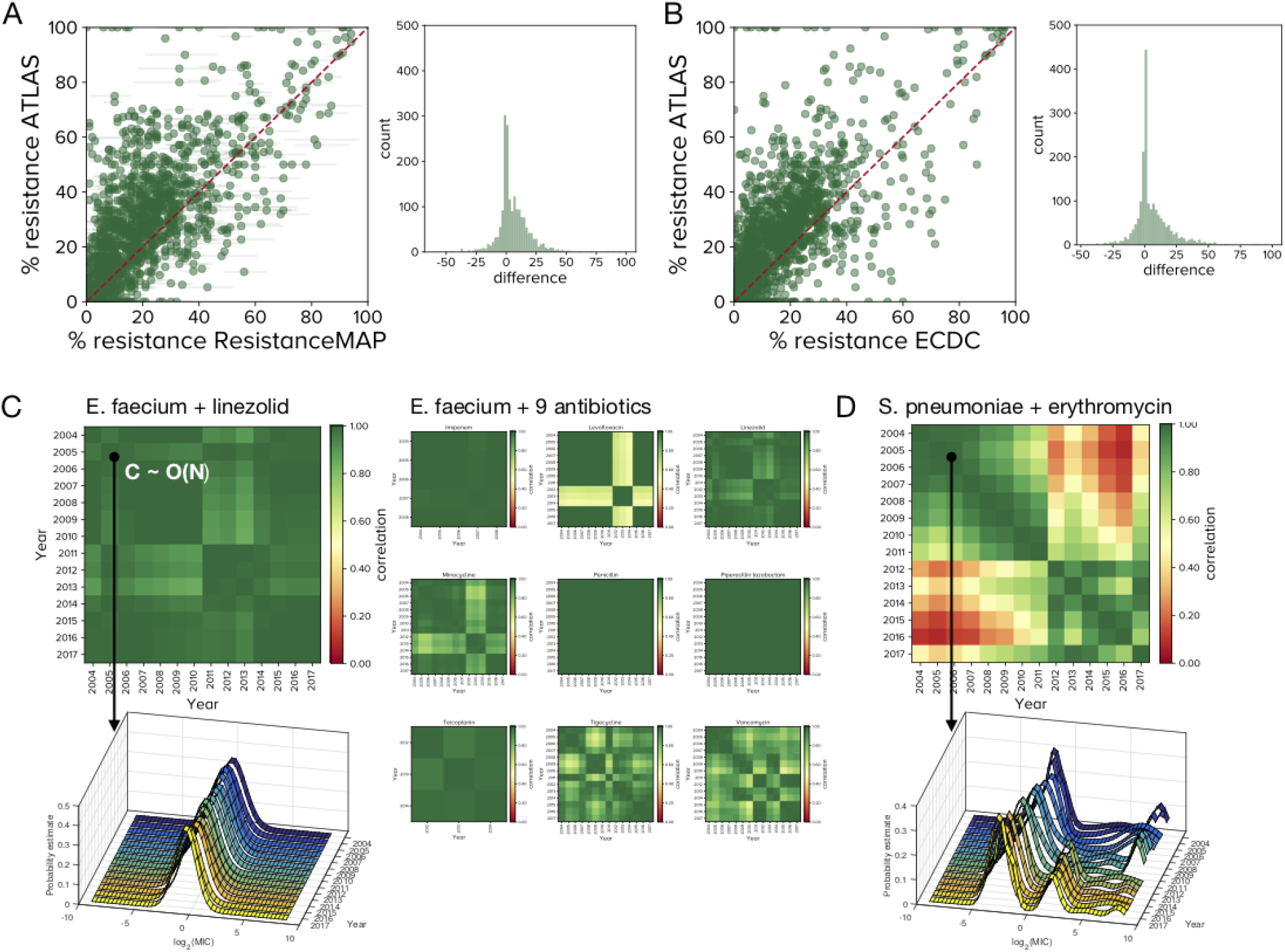
Data quality within and between-databases. Atlas over-estimates the frequency of resistance relative to A) ECDC and B) Resistance Map databases. Histograms of differences between the %age of resistant cases in Atlas and the other databases are positively skewed (*p* < 0.001) with a mean difference of approximately 10%. C) Between-year correlations show Atlas’s PA pairs are skewed towards unity (here green; Figure S5C, S6). The left column shows the global MIC distribution of *Enterococcus faecium* and linezolid is stable and the correlolgram is close to the unity matrix. The middle panel shows 9 other antibiotics’ yearly correlolgrams, a banded structure indicating highly correlated MIC distributions with larger changes in some years. D) *Streptococcus pneumoniae* and erythromycin have a pronounced 2-cluster correlelogram structure because the MIC distributions correlates poorly between some years: a high-MIC cluster diminishes and is replaced by a cluster with lower MIC in 2012. The main text hypothesises reasons for this because this is replicated with clindamycin (Figure S13).

### Within-database consistency: Atlas year-on-year

Being the result of a clinical assay, each Atlas datum is not replicated nor has it any intrinsic measure of uncertainty. It is therefore plausible that MICs of a given PA-pair exhibit no between-patient correlation or between-year coherency and, rather, resemble a noise process. As MICs are resolved only by year, each PA pair does not yield an MIC timeseries but rather a set of MICs of different cardinalities for each year. So, we asked whether empirical MIC distributions from year *y_i_* would correlate with year *y_j_*; we call the symmetric matrix of all year-year correlations, *C_ij_*, a ‘correlelogram’. Figure 1C provides an exemplar correlelogram for *Enterococcus faecium* and linezolid and correlelograms for 9 other antibiotics. A test of the distribution of singular values of *C_ij_* shows Atlas has significantly greater correlations between-years for each PA pair than expected for noise processes (Figure S5) and PA pairs with the lowest between-year correlations typically have data for the fewest years (Figures S6). Thus MIC datastreams of sufficient duration can form coherent, if slowly changing, datasets for several years (Figure S7).

That said, some MIC distributions change abruptly. For instance, *Streptococcus pneumoniae* and erythromycin have a modular correlelogram structure (Figure 1D) reflecting the seeming loss of a high-MIC subpopulation in 2011 and this illustrates how Atlas can capture prevalent clinical conditions. Inspecting this PA pair more closely shows international definitions of susceptibility for *S. pneumoniae* were revised in 2008 resulting in more strains appearing susceptible^*27*^ but this change would not affect raw MICs so it cannot explain Figure 1D. However, the introduction of conjugate vaccines (7-valent in 2000, 13-valent in 2010) reduced the incidence of antibiotic resistant invasive clones and, in particular, clindamycin resistance fell following the 2010 vaccine in developed countries.^*28,29*^ As *erm* genes confer resistance to both erythromycin and clindamycin^*30,31*^ Figure 1D may reflect a correlated change in high-level resistance to both antibiotics and, indeed, we observe very similar dynamics in the analogous correlelogram for clindamycin that also exhibits a marked shift in 2011 (Figure S13).

### Reconciling quantitative genetics with the clinical view: cluster of greatest MIC (*R*)

That Atlas is not an ensemble of uncorrelated MICs, though nor is it a set of stationary MIC distributions, are consistent with evolutionary dynamics acting on the MIC as a phenotype. A standard quantitative genetics approach to elucidating those dynamics would be to linearly regress MIC against time which we did for every PA pair, first globally (Figure 2A-B) and for Europe (Figure S8A). Figure 2A summarises the predicted changes and a curious feature is apparent: apart from South-East Asia and Central America, particularly India, China and also Ireland, Serbia and Croatia, more global mean MIC reductions are observed than increases from 2005-2015 (skewness test, *p* < 0.001) whether, or not, US (whose data dominates Atlas, Figure S2) is excluded from the analysis. Given reports of increased resistance and the fact that Atlas exhibits typically greater frequencies of resistance than ECDC and ResistanceMap data (Figure 1A-B), this seems anomalous. To explain this, it could indicate MIC distributions where the highly resistant tail has different dynamical behaviour to the mean, so we tested this idea.

**Fig. 2:**
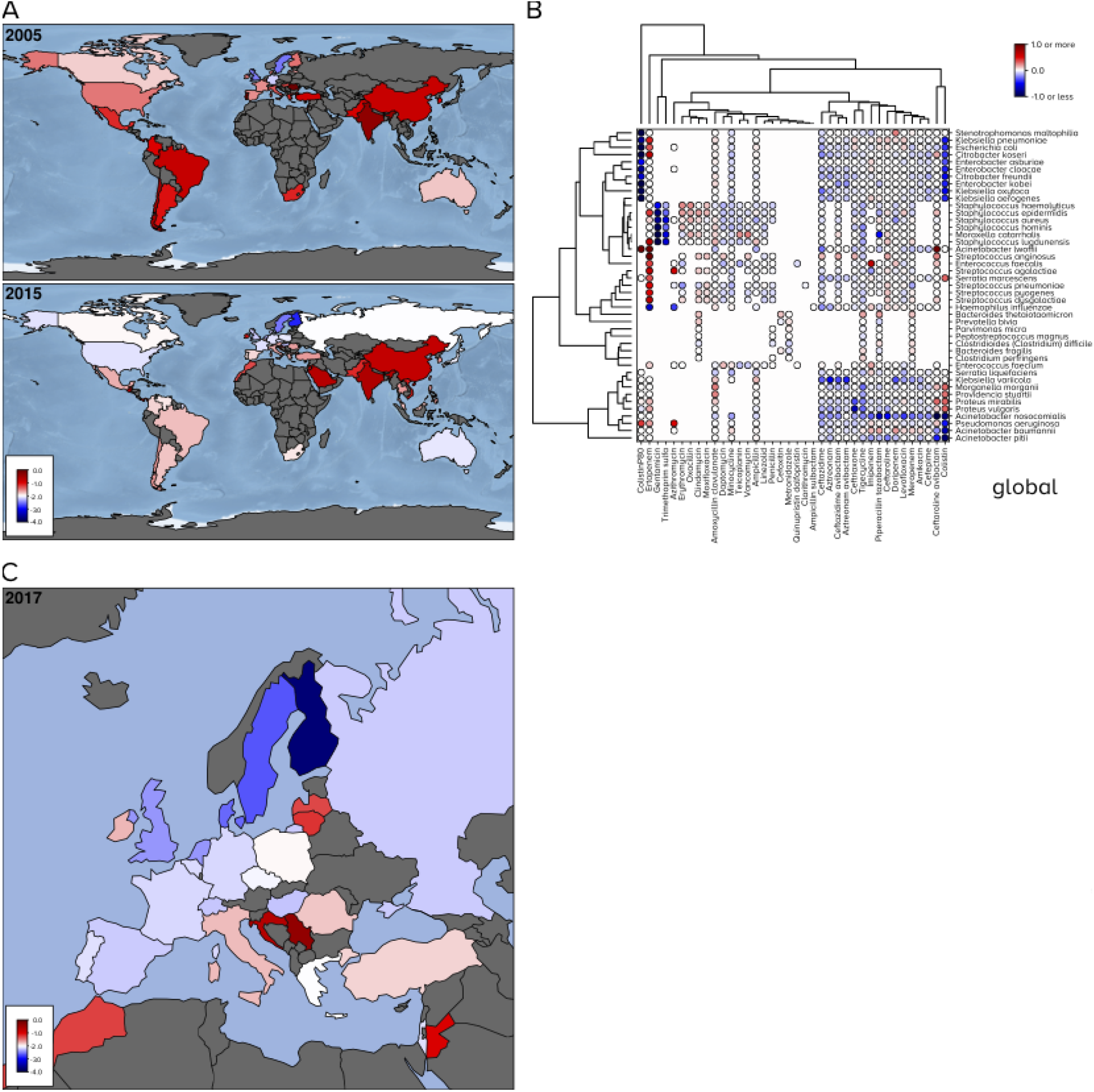
Dynamics of mean MICs: some decreases with increases in Asia. Linear regressions predicted MIC changes for all PA pairs: white values show no available data for a PA pair, a ringed circle on a blue-red colourscale indicates predicted change (red is +ve log_2_ MIC change per year, blue is −ve, grey no change). A) The global distribution of mean MIC aggregated across all PA pairs: possible reductions form 2005-2015 in some countries with increases in India and China. B) Global mean MICs resolved by PA pair are clustered into pairs of similar change indicative of a global motif of increases and decreases. The analogous figure for Europe is similar (Figure S8A). C) Mean European MICs aggregated across all PA pairs in 2017 shown by country indicate differences between eastern and western Europe.

Linear regression is too a coarse method for that, so we used a clustering methodology to extend the clinical categorisation into susceptible (S) and resistant (R) strains by, instead, seeking subpopulations with the highest and lower MICs. By modelling each MIC distribution as a Gaussian mixture, a superposition of *k* different normal distributions, we determined an information-optimal *k* for each PA pair for each year. The sub-population R was then defined as the cluster with greatest mean MIC and S was defined as the complement of that; the probabilisitic boundary between S and R is the MIC value with an equal likelihood of being in either. For robustness, we repeated this methodology on a library of 50 synthetic replicates of Atlas, each with small-variance noise added to every MIC (Methods). To determine MIC changes, we applied longitudinal regression to S and R separately to each Atlas replicate (Figure 3A) and this predicts the changes of R are increasing, and *not* decreasing, for many PA pairs (Figure 3B and D). The above anomaly arises, therefore, because many PA pairs have mean MIC decreases and yet their R-MIC increases (Figure 3C).

**Fig. 3:**
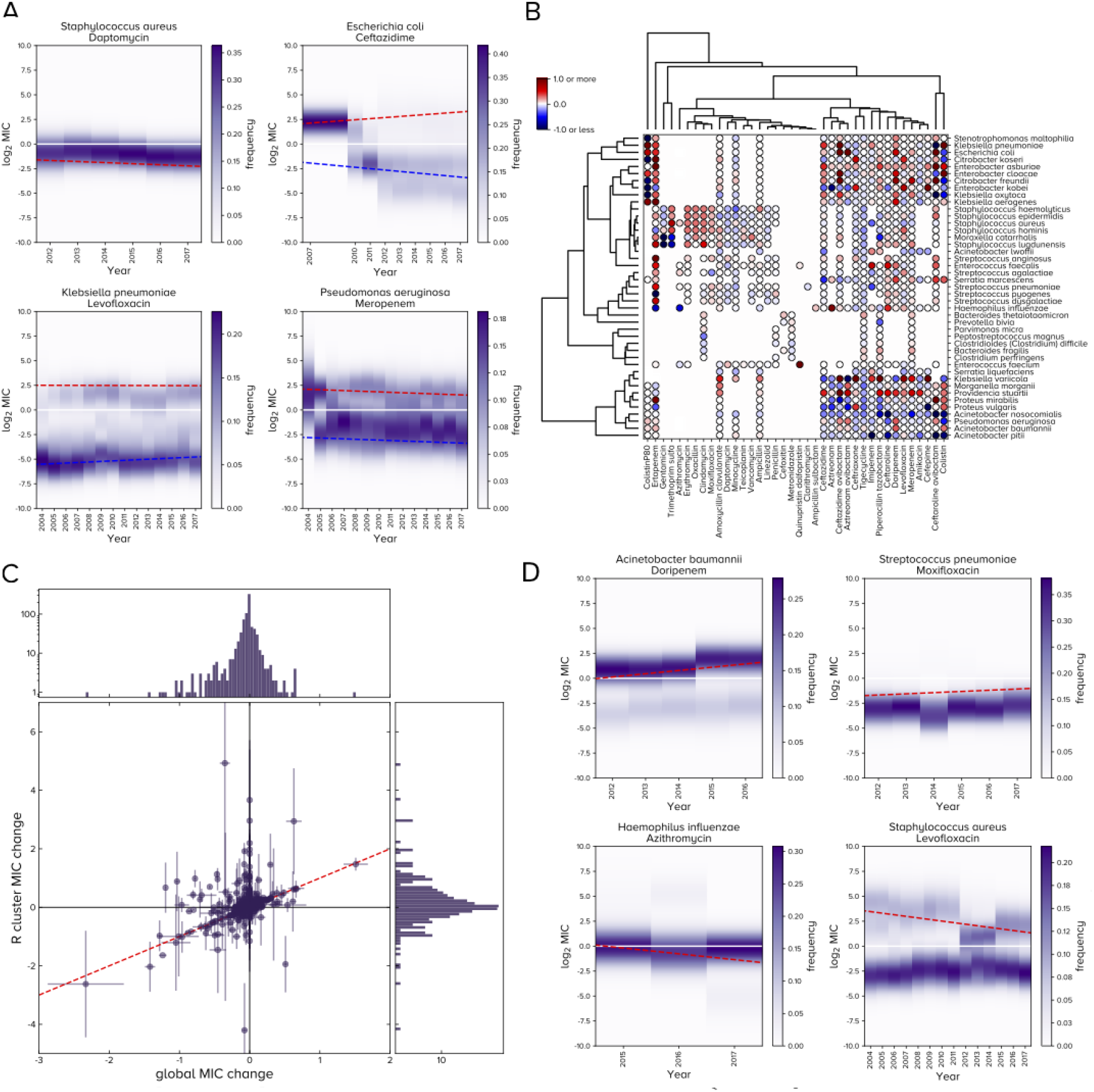
Clinically relevant MIC dynamics. A) Longitudinal changes in mean S and R MICs are shown for 4 PA pairs: these exemplars have 1, 2 or 3 clusters each and linear regressions capture MIC changes of R (red) and S (blue). B) The latter MIC changes are shown for each PA pair on a red (+ve) to blue (-ve) scale in a clustergram on a scale of log_2_ MIC per year for global data; the analogous plot for European data is similar (Figure S8B). C) Data either side of the line ‘*y* = *x*’ (red dashes) show mean MICs of R and population mean MICs are positively correlated in some cases (*ρ* = 0.33) but there are also many PA combinations where R decreases in MIC and yet population mean MIC increases are predicted; this is evidence of disruptive selection. (Compare C with Figure 2B and 3B (or with Figures S8A and B for Europe-only data) which also illustrate this.) D) These exemplars illustrate each of 4 possibilities: PA pairs whose R cluster is either sub- or super-clinical, where its MIC is either increasing or decreasing.

This observation has an evolutionary explanation from population genetics. Our prior expectation was that S and R would increase in their respective mean MICs for all PA pairs but cases where S is stable or decreasing while R-MIC increases (Figure 3C) may indicate ‘disruptive selection’. This arises when a population diverges genomically, possibly as sub-populations respond to different selection pressures. Figure 3C indicates putative cases of ‘purifying selection’ too where the MICs of S and R tend towards a common value and Figure 3D has the most optimistic case of all whereby S is stable, or decreasing marginally, while the R-MIC is decreasing. Disruptive selection is the most common of these cases and R MIC increases are typically greater than the respective global mean change (Figure 3C).

### Clinical Observations

To detail these dynamics we introduce two phase planes: first, a 2-dimensional depiction of (*R, dR/dt*) (Figure 4A) and, second, a plot of (*dS/dt, dR/dt*) (Figure S11). The latter is consistent with disruptive, purifying and directional selection (Figure S11) but the former is more useful: it highlights where the most clinically relevant cluster, R, is now and where it is heading. Figure 4B summarises (*R, dR/dt*) phase planes of the entire Atlas database as one heatmap and, to illustrate how the phase planes are determined, Figure 3D illustrates exemplar R clusters and their MIC dynamics. Phase planes of 8 clinically important pathogens (Figure 4) visually skew towards MIC increases and while Atlas has more PA pairs than are in current clinical use, all data derive from clinical tests. Figure 4 summarises some clinically relevant trends consistent with previous reports but also others that are less expected, as follows.

**Fig. 4:**
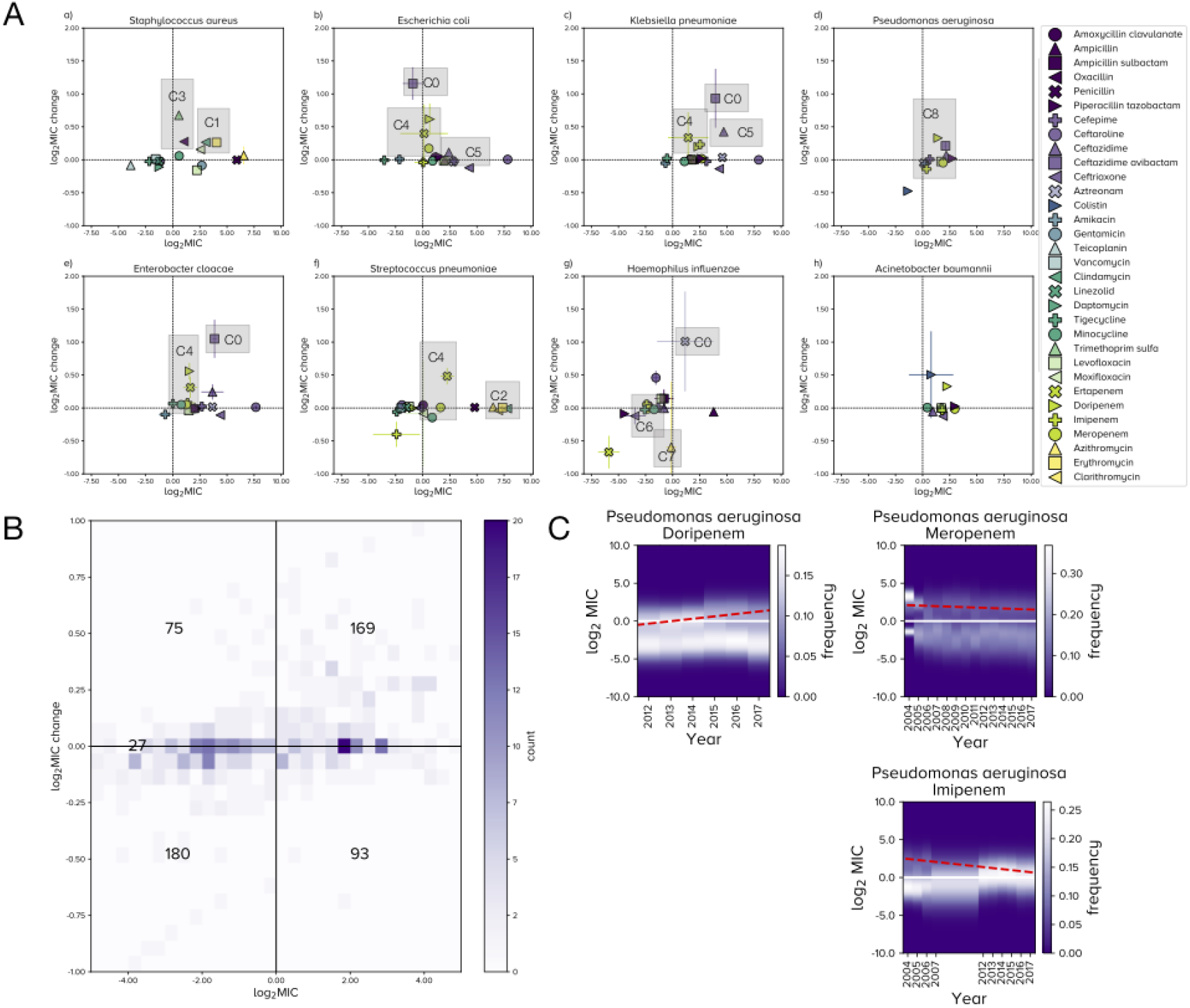
Eight clinically important pathogens and their R derivatives. A) Each panel is a phase plane showing (*R,dR/dt*) (± 1 synthetic standard deviation, *n* = 50, see Methods) where the x-axis shows the mean MIC of R and *dR/dt* is its yearly derivative. *Acinetobacter baumannii* exhibits clinical resistance to all these antibiotics and most phase planes are skewed towards resistance. Labels C0-C8 are clinically relevant observations referred to in the text. B) A 2d histogram shows where 544 PA pairs are situated in the (*R,dR/dt*) phase plane (note: 544 out of 827 studied PA pairs have a clinically defined breakpoint, so we can only use those in (*R,dR*) phase planes). C) Changes in the mean MIC of R of global data for *Pseudomonas aeruginosa* and 3 carbapenems: doripenem resistance is increasing from sub- to super-clinical whereas meropenem appears stable. The R cluster of imipenem is slowly decreasing according to a regression (red dashes) but, in fact, this has merged with the S sub-population as it achieves super-clinical dosages.

As expected, *Acinetobacter baumannii* exhibits resistance to all antibiotic classes with scant evidence of MIC reductions (Figure 4A) and reports of colistin resistance in *A. baumannii*^*32*^ are consistent with an R cluster that we first observe in Atlas in 2015 (Figure S9(b)). The most rapid MIC increases in Atlas are for ceftazidime avibactam (CAZ-AVI; Figure 4A, label C0) for instance, an emerging R-cluster first detected for *Klebsiella pneumoniae* in 2014 is increasing in MIC (Figure S9(a)). Other enterobacteriaceae (*E. coli* and *E. cloacae*) also have a large change in R-MIC for CAZ-AVI (Figure 4A, label C0) because an R cluster recently appeared. However, clinical testing behaviour means that MICs of such second-line agents must be interpreted with care. CAZ-AVI is often tested against strains resistant to frontline treatments^*33*^ so the likelihood of reporting as resistant may be biased. Thus, when its MICs are contrasted with earlier surveys conducted to determine its wider efficacy, this could look to Atlas like a resistance shift that merely reflects that bias and this issue may affect the newest antibiotics most. A large change in R-MIC is also predicted for *H. influenzae* and aztreonam (Figure 4A, label C0) but with large uncertainty estimates under synthetic Atlas replication (Methods).

As with *S. pneumoniae*, erythromycin and clindamycin data exhibit similar behaviour against *S. aureus* but in this case resistance to both is increasing at similar rates (Figure 4, label C1), again consistent with *erm genes*.^*30,31*^ Conversely, and consistent with reports of a plateau,^*34*^ erythromycin resistance in *S. pneumoniae* is high but not increasing (Figure 4, label C2) and its MIC distributions support this observation (Figures 1D, S13).

*S. aureus* resistance to trimethoprim sulfamethoxazole (TMP-SMX) and oxacillin are both increasing (Figure 4, label C3). Oxacillin is a penicillin that has substituted methicillin, TMP-SMX is used against methicillin-(and thus oxacillin-) resistant *S. aureus*^*35*^ and this trend suggests TMP-SMX is failing to mitigate resistance in *S. aureus*. Secondary prescribing behaviour may be important here as methicillin-resistant *S. aureus* clones emerged from antibiotic-susceptible community lineages and TMP-SMX became an important oral agent for community-acquired non-multiresistant MRSA (cMRSA) which may have driven TMP-SMX resistance. The R-MIC of TMP-SMX is significantly above the clinical breakpoint which implies the treatment of MRSA using TMP-SMX may be at risk.

In terms of beta lactams, Atlas exhibits the following. Carbapenem MICs for *K. pneumoniae* have been reported as bimodally distributed where high MICs reflect outer membrane protein (OmpK36) defects,^*36–38*^ this is consistent with long-term distinct S and R clusters in Atlas (Figure S10). Carbapenem resistance is increasing^*39*^ among enterobacteriaceae and Atlas reflects this (Figure 4A, label C4; *E. coli, K. pneumoniae, E. cloacae*, also *S. pneumoniae*). Porin defects in *K. pneumoniae* have poor ertapenem uptake^*40*^ and we hypothesis porin mutants explain increasing ertapenem MICs in *E. cloacae*^*41*^ (Figure 4A, label C4). Ceftazidime is following prior trends in uropathogenic *E. coli*^*42*^ that derive from mobile CTX-M-type genes^*43*^ that are dominated by a ceftazidime-resistant phenotype. These may contribute to a rise of ertapenem resistance in *E. coli* and *K. pneumoniae*.^*44*^ The rate of ceftazidime MIC increases for *K. pneumoniae* are greater than *E. coli* (Figure 4A, label C5) which might be explained by the former evolving carbapenem resistance through porin mutations^*40*^ thus increasing their frequency in bacterial populations that share mobile beta-lactamase genes among enterobacteriaceae.

In contrast with smaller studies^*45,46*^ where differences in carbapenem resistance were not detected, the R MIC of doripenem is increasing more quickly in Atlas than other carbapenems against *P. aeruginosa* (Figure 4A, label C8). This may be due to changes in efflux-mediated cross-resistance between carbapenems,^*47*^ more speculatively it might even have resulted from a change in manufacturing or usage base as carbapenem patents expired in the decade after 2010. Seeking to better understand this, we found doripenem has the fastest increasing carbapenem R MIC in almost all countries (Figures S12) thus between-country differences cannot explain doripenem’s rise. S- and R-MICs of doripenem against *P. aeruginosa* are converging towards those of meropenem (Figure 4C, year 2017) so we speculate doripenem’s increase might result from recommendations shifting to mitigate resistance in other carbapenems.^*48–50*^ Consistent with this, a 2019 survey of 20 US hospitals shows while doripenem has more uncertain usage data, it was the most used carbapenem when measured in days of treatment per patient day (DOT-PPD) whose defined daily dosage (DDD-PPD) was approximately half of the most-used carbapenems (51, Tables 1 and 3).

*H. influenzae* is susceptible to minocycline with no significant increase in R-MIC (Figure 4A, label C6) so we predict this can be used against beta lactam-resistant *H. influenzae*. We find no evidence of azithromycin resistance in *H. influenzae* and, interestingly, resistance may be marginally decreasing (Figure 4A, label C7). However, the number of cases supporting this observation is fewer than 100 per year in a narrow year range (2015 to 2017) and the uncertainty measure from synthetic Atlas replication is not significant.

## Discussion

Our methodology supplements Atlas with metadata that can be queried to detect PA pairs with important properties. We have sought problematic PA pairs where the R-MIC is super-clinical and still increasing: *Acinetobacter baumannii* and doripenem is one such pair (Figure 3D). Conversely, we can seek PA pairs where, importantly, the R-MIC is subclinical but increasing, like *Streptococcus pneumoniae* and moxifloxacin (Figure 3D). Seeking some optimism, we can also seek pairs where the R-MIC is subclinical and decreasing, like *H. influenzae* and azithromycin, and also ones where the R-MIC is super-clinical but decreasing, like *S. aureus* and levofloxacin (Figure 3D). Of course, other PA pairs exhibit similar behaviour and this demonstrates the need to analyse subpopulations, not just frequencies of super-clinical MICs, to better understand resistance. Having done so, the resulting patterns appear as complex as might be expected in a large, spatially extended evolutionary ecosystem that hosts bacteria.

We, inevitably, have concerns that Atlas contains biases and inconsistencies that we cannot detect algorithmically or rationalise clinically, given the sheer volume of data, but we have also found that Atlas exhibits many signals consistent with prior clinical expectations. Given this, we made the working assumption that Atlas data may be least reliable among the PA pairs whose year-year MIC correlations are lowest (Figure S6). So with this caveat, if Atlas curators continue to grow their database, it must surely prove an invaluable resource for understanding clinically relevant changes in MIC signals that might not be detected in smaller cohorts, or even in the data of single countries. If Atlas can be supplemented with more fine-grained metadata on antibiotic consumption, we might be able to discern how increasingly rational antibiotic usage mediates resistance on large spatial scales.

## Methods

Atlas data have the following structure: one record represents one patient and comes with metadata, including the year and country of infection, the isolated pathogen strain, whether the latter was deemed resistant, susceptible or intermediate against a panel of antibiotics and it lists all numerical MICs used for those classifications. Data amalgamate surveillance programmes across approximately 70 countries so there is no uniformity between records. Thus, each MIC is presented un-replicated, with no measure of variation, where the panel of antibiotics assayed vary between patients, even for the same pathogen. MICs are quoted as scaled powers of 2, *A* · 2^−*d*^ say, where *A* is a baseline dose quoted in *μg/mL* and typically *d* is the critical two-fold antibiotic dilution found using an MIC-determination protocol. Other laboratory protocols provide MICs too, like disk diffusion tests. To ensure uniformity, we find MIC breakpoints by determining the maximal dose in the database deemed ‘susceptible’ for each PA pair and we fix this value. We then divide each MIC by this breakpoint, thus scaling out the dose constant A, and we log_2_-transform the result. Physical units for the derived MIC are the numbers of 2-fold dilutions relative to the breakpoint, so a value of ‘MIC = 0’ represents ‘clinically resistant’ throughout. Throughout, wherever the term ‘MIC’ is used, including all figures, it always refers to the log_2_ value relative to a clinical breakpoint, as described in the main text, raw database MIC values are never shown.

Computations were performed using Pandas in Python 3 and Matlab 2018, including the Statistics Toolbox. Atlas data is publicly available following website registration*, data and further information can be downloaded from the following links:

project overview: https://amr.theodi.org/project-overview
project description: https://wellcome.ac.uk/sites/default/files/antimicrobial-resistance-surveillance-sharing-industry-data.pdf
data download*: https://www.synapse.org/#!Synapse:syn17009517/wiki/585653, and also the following link: https://s3-eu-west-1.amazonaws.com/amr-prototype-data/Open+Atlas_Reuse_Data.xlsx
All analysis codescan be downloaded here: https://github.com/PabloCatalan/atlas

### MIC variation: additive noise model

Raw Atlas MICs are discrete data quoted without uncertainty measures or replication, to test the robustness of our analyses we created 50 synthetic replicates of Atlas data to form uncertainty estimates, as follows. We simulated within-replicate MIC variation of the kind generated by laboratory assays by adding a stochastic quantity to each MIC sampled from a normal distribution with a variance of 1 dilution. This approach was taken to perform a sensitivity analysis where we tested which patterns of resistance and resistance change were robust and significant given this degree of stochastic noise in MIC and which were not.

### Determination of S and R and their derivatives

We used Gaussian mixture models to determine the optimal number of clusters in the distribution of each MICs for each PA pair and these models identified the cluster with greatest mean MIC; this is called the R-cluster, or just R. The boundary of the R cluster was defined using a fixed confidence threshold and all MICs below this were labelled as the S subpopulation. Linear regressions were then used to determine MIC change versus time for S and R (Figures 3A/D and 4C).

## Acknowledgements

We thank the AMR Research Initiative and the Open Data Institute for publishing their dataset, free of access charges. ResistanceMap data was very kindly supplied by Quentin Leclerc to whom we express our thanks, we also thank Github for hosting all the codes and open data associate with them.

## Data availability

All data is publicly and freely available, it can be downloaded from relevant websites at no charge and codes are hosted on Github, see Methods.

## Author contributions

All co-authors interpreted data and, in addition, PC provided funding, designed and optimised algorithms, analysed data, wrote and edited the manuscript; CR suggested analyses and edited the manuscript; JB suggested analyses and edited the manuscript; IG provided funding, suggested analyses and edited the manuscript; JI suggested analyses, wrote and edited the manuscript; RB designed and optimised algorithms, analysed data, wrote and edited the manuscript

## Supplementary Information

### Changepoint analysis for low-MIC sampling bias in Atlas

Given a time series consisting of a number of MICs, *n*_1_,…, *n_T_* say, we want to test the null hypothesis, *H*_0_, that all sampled points come from the same distribution against the alternative hypothesis, *H*_1_, that there is a single changepoint in mean, *μ*, at timepoint *τ*:

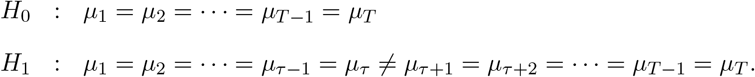

Assuming that the sample is normally distributed and that the variance, *σ*^2^, does not change across the sample, the log-likelihood ratio between the two hypotheses is:

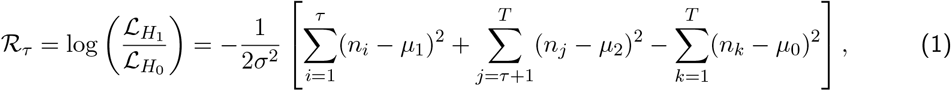

where 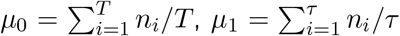 and 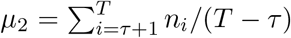. We then find the value of *τ* that maximizes 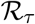, and we define

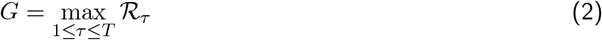

where 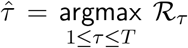. We accept the existence of a changepoint, at time 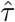, if *G* is larger than a critical value, λ^*^, defined by the Bayesian Information Criterion^*52*^ 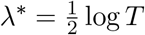.

**Fig. S1:**
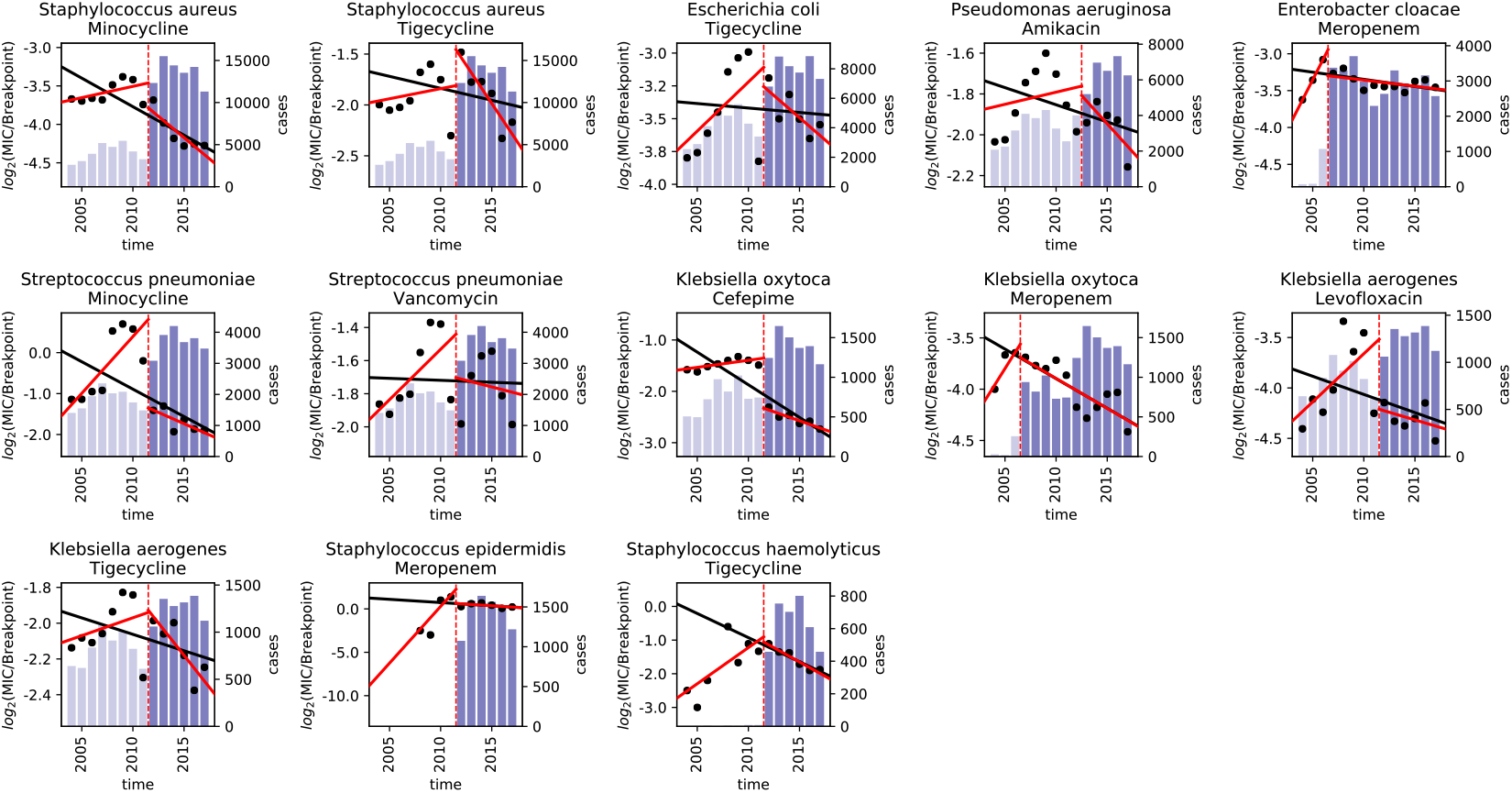
A changepoint analysis detects PA pairs for which temporal decreases in MIC coincide with increased antibiotic susceptibility testing. Each sub-plot shows analogous data: first, the blue bars represent the yearly global number of clinical tests for each PA pair and the change from light to dark blue highlights the year at which a change in this number was detected using a changepoint analysis. The two red lines show slopes of linear regressions that indicate significant increases in MIC before the changepoint (the vertical dashed red line), followed by a significant decrease after the changepoint. Finally, the black line is the result of applying a regression to the entire MIC timeseries to estimate a coarse rate of change, this is decreasing with time in all cases. Thus, these PA pairs represent biases whereby predicted mean MIC decreases may be due to an increase in clinical testing of less resistant pathogens through time.

**Fig. S2:**
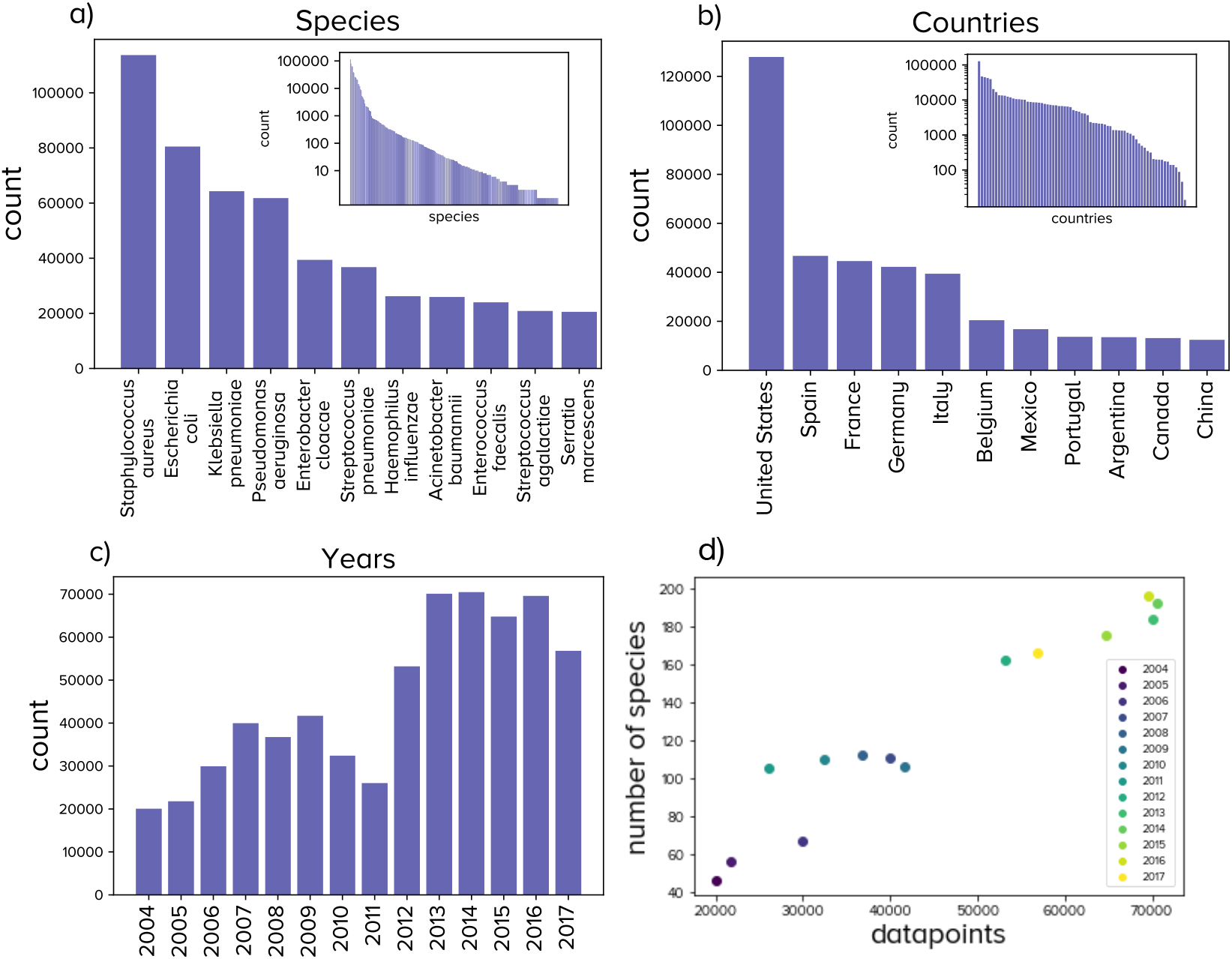
Basic statistics of the Atlas data. a) The bacterial species most represented in Atlas. b) Some countries contribute more data than others, even after accounting for population size. c) More MIC data has been added to the Atlas database in recent times with a noticeable increase around 2012; this may be due to more countries having more data to contribute to the programme through time. d) An analogous comment applies to bacterial species that increase in number through time.

**Fig. S3:**
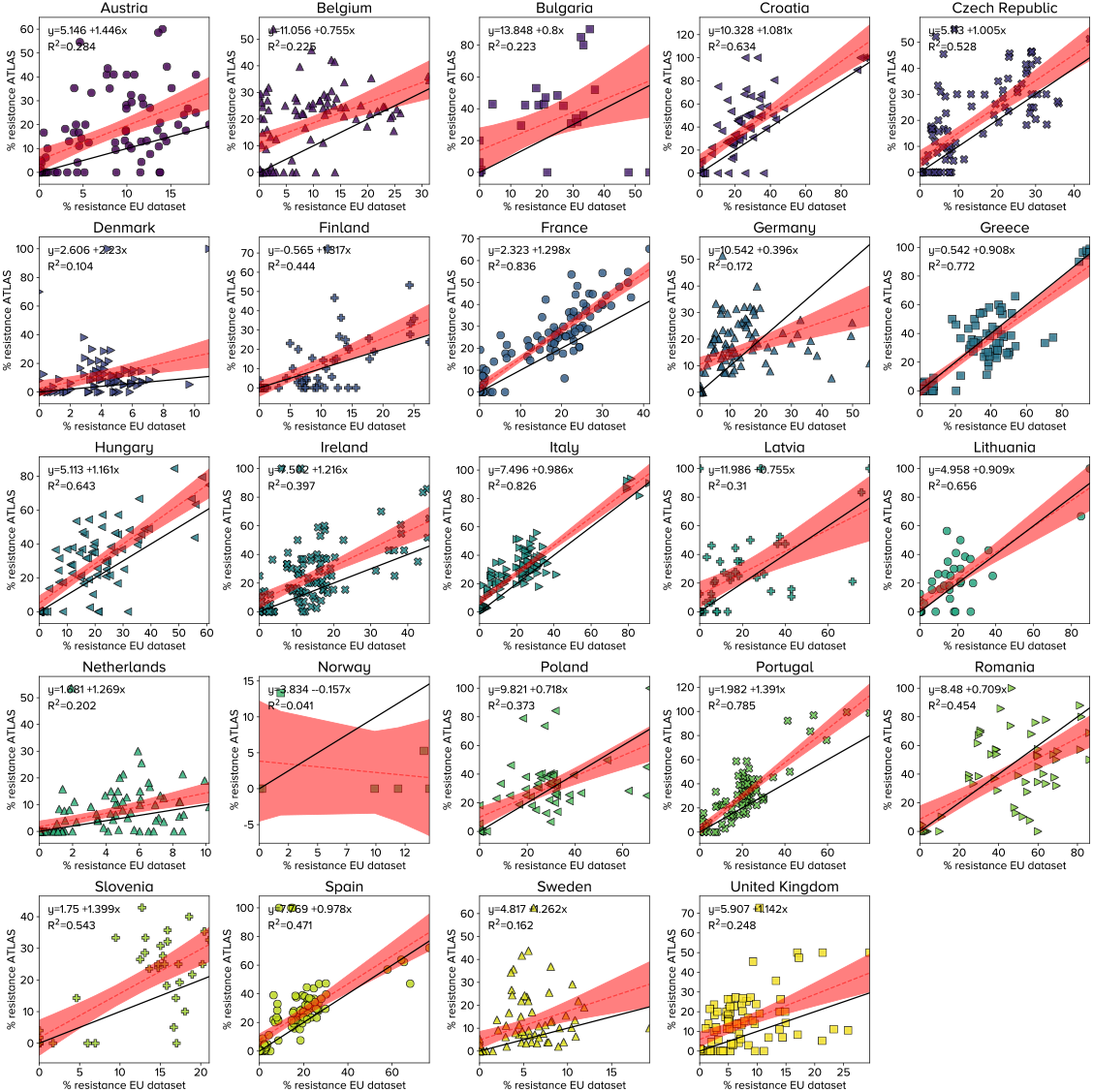
Collated ECDC and Atlas data and their correlations for European countries. Linear regressions indicate Atlas reports higher frequencies of resistance than ECDC data but only in some countries (each symbol denotes one PA pair). The linear regressions, which should report *y* = *x* can yield *y* = *ax* + *b* where (*a, b*) is significantly different (1,0) in several cases.

**Fig. S4:**
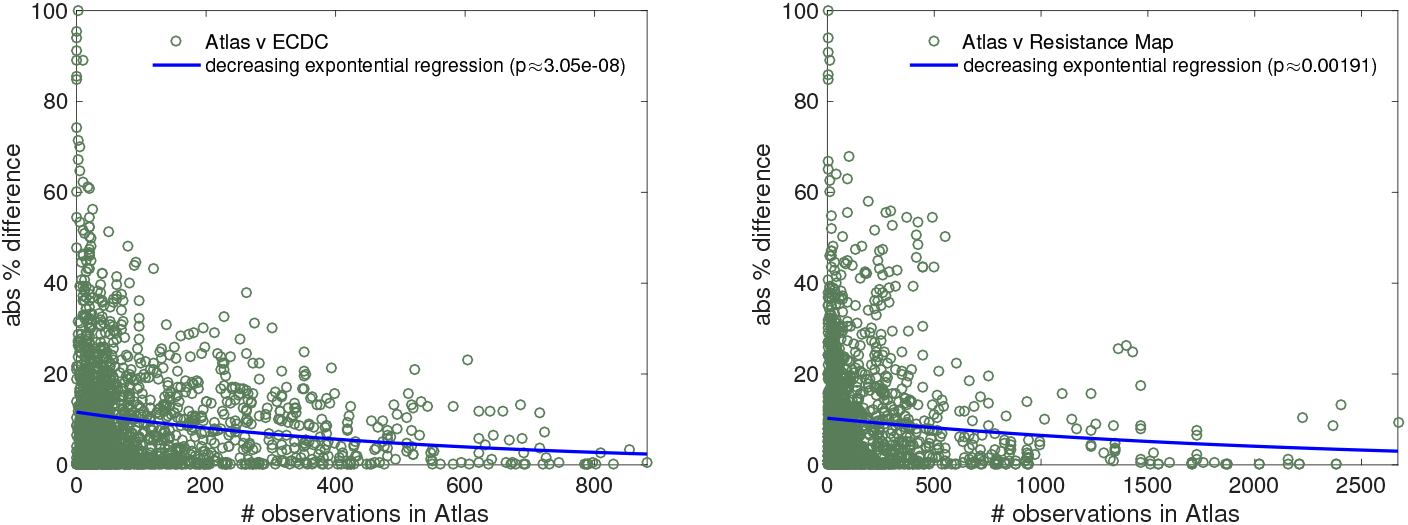
Comparing % resistance of PA pairs in each country to Atlas from. **a) ECDC and b) RMap databases:** larger within-country datasets per PA pair have smaller between-database differences (exponential regression *p* values in legend and fits to data in blue).

### Spectral Analysis of Correlation Matrices

Given the lack of replication in clinical MIC assays, it is important to ask whether MIC timeseries from the Atlas database are noise-like or are they formed from coherent signals drawn from multiple patients with small noise? We therefore use a test for MIC coherency between years using the following null model: we ask what within- and between-year correlations would be found in MICs if they were drawn from uncorrelated processes each year and then test the Frobenius norm of the correlation matrix of each PA pair defined in the main text for consistency with that null model.

So, assume a data stream, (*t_k_, m_k_*), of MIC values (at time *t_k_*) is taken for *N* years for a given PA combination, where *i* and *j* are labels for two years and where each MIC is a random variable with *m_k_* ~ PDF(*f*) where *f* is a probability density function that does not depend on year and *F* is its cumulative distribution function supported on the discrete set {–10, –9, –8,…, 8, 9,10} =: *M*, where ±10 is a clinically reasonable upper and lower bound for MIC observations on the log_2_ scale as defined in Methods.

Now bin data according to the year associated with *t_k_* and let *y_j_* be the sampled MIC distribution for year *j*, and so we expect *y_j_* ≈ *f* as a vector supported on *M* (of dimension 21, the cardinality of *M*). Centralise the sample distribution data, *Y_j_* = (*y_j_* – mean(*y_j_*))/std(*y_j_*), so each *Y_j_* is a random variable with some distribution function *Y_j_* ~ PDF(*g*) where *g* derives from *f* and is positive, takes on continuous values in [0,1] and is supported on 2*M*. Now set *ŷ_ij_*:= *E*(*Y_i_* – *Y_j_*)^2^ = 2(1 – *E*(*Y_i_Y_j_*)) and let *X* = (1 – *ŷ_ij_*/2) be a correlation matrix which is symmetric with positive entries between 0 and 1. Moreover *X* can be written *X* = *I_N_* + *L* + *L^T^* where *L* is a lower diagonal matrix with entries between 0 and 1 where *I_N_* is the identity matrix.

If yearly MIC data is correlated because MICs can be modelled as a stationary random variable with the same distribution each year then *Y_i_* and *Y_j_* both have *g* as their PDF. Thus 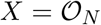, which is the *N* × *N* rank-1 matrix consisting only of 1 in each of its *ij* entries. On the other hand, if yearly MICs have been drawn each year from independent random variables then *E*(*Y_i_Y_j_*) = 0 and *X* = *I_N_*, the *N* × *N* identity matrix.

Intuitively, if all MICs of a PA pair are drawn from a stationary MIC distribution then *X* will be a matrix of rank 1. If MICs are drawn from non-stationary distributions of MICs subject to evolutionary change because bacteria are evolving in response to antibiotic use, we should expect some changes in between-year MIC distributions, so *X* will have rank greater than 1. In the extreme case that uncertainty from clinical procedures concerning MIC measurements, and the fact that different assays are used, renders the MIC a noise-dominated process then *X* should have rank *N*.

We now test these algebraic properties of *X* and the Frobenius norm of *X*, ║*X*║_*F*_ = tr(*X^T^X*)^1/2^, is helpful in this regard. The eigenvalues of *X* are counted according to algebraic multiplicity for the following eigenvalue sums. So, if *σ*(*A*) denotes the spectrum of a matrix, *A*, then yearly MICs are uncorrelated when *σ*(*X*) = {1,1,…, 1} and rank(*X*) = *N* but they are correlated when *σ*(*X*) = {1, 0,…, 0} and rank(*X*) = 1. Now singular values of *X, s*(*X*), satisfy *s*(*X*) = *σ*(*X^T^X*) = *σ*(*X*)^2^ and

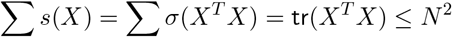

because each element of *X* can be bounded by the Cauchy-Schwartz inequality under assumptions of the null, where tr denotes the trace of a matrix and tr(*X^T^X*) is the sum of squares of all the entries in *X*. Also

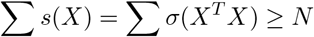

because the *N* diagonal entries of *X* are all 1.

Given this, we now define functionals *τ* and *τ_N_* that can be used as the basis of a test that compares correlation matrices for PA pairs independently of the value of *N* over which the MIC data stream has been gathered. As *X* = *L* + *I* + *L^T^*, where *L* is a lower-diagonal matrix with entries between 0 and 1, then we are interested in whether the mean square off-diagonal entry in *L*, *l_ij_*, is 0, 1, or neither. Now, this mean square, noting tr(*X*) = tr(*I*) = *N*, is

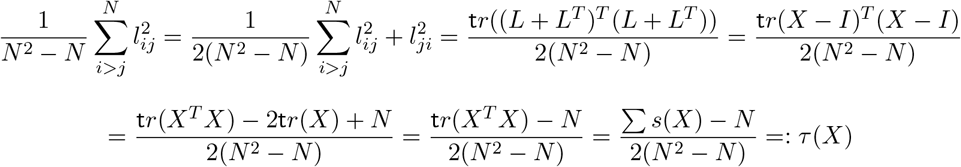

where *τ* lies between 0 and 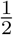.

Or, for a fixed N, we can test the total lower diagonal diagonal square entries of *L*, namely

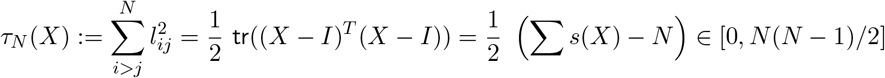

for a single PA pair, as follows. A PA pair has stationary yearly MICs if tr_*N*_(*X^T^X*) = *N*(*N* – 1)/2 and a PA pair has uncorrelated yearly MICs if tr_*N*_(*X^T^X*) = 0. In practise, Figure S5 shows that histograms of *τ_n_* for *N*-year timeseries are closer to *N*(*N* – 1)/2 than to 0, consistent with non-stationary but positively correlated year-to-year MIC distributions despite all MICs being drawn from different patients. Similarly, the histogram of *τ* across Atlas is biased away from zero (Figure S5b, *p* < 10^-15^).

**Fig. S5:**
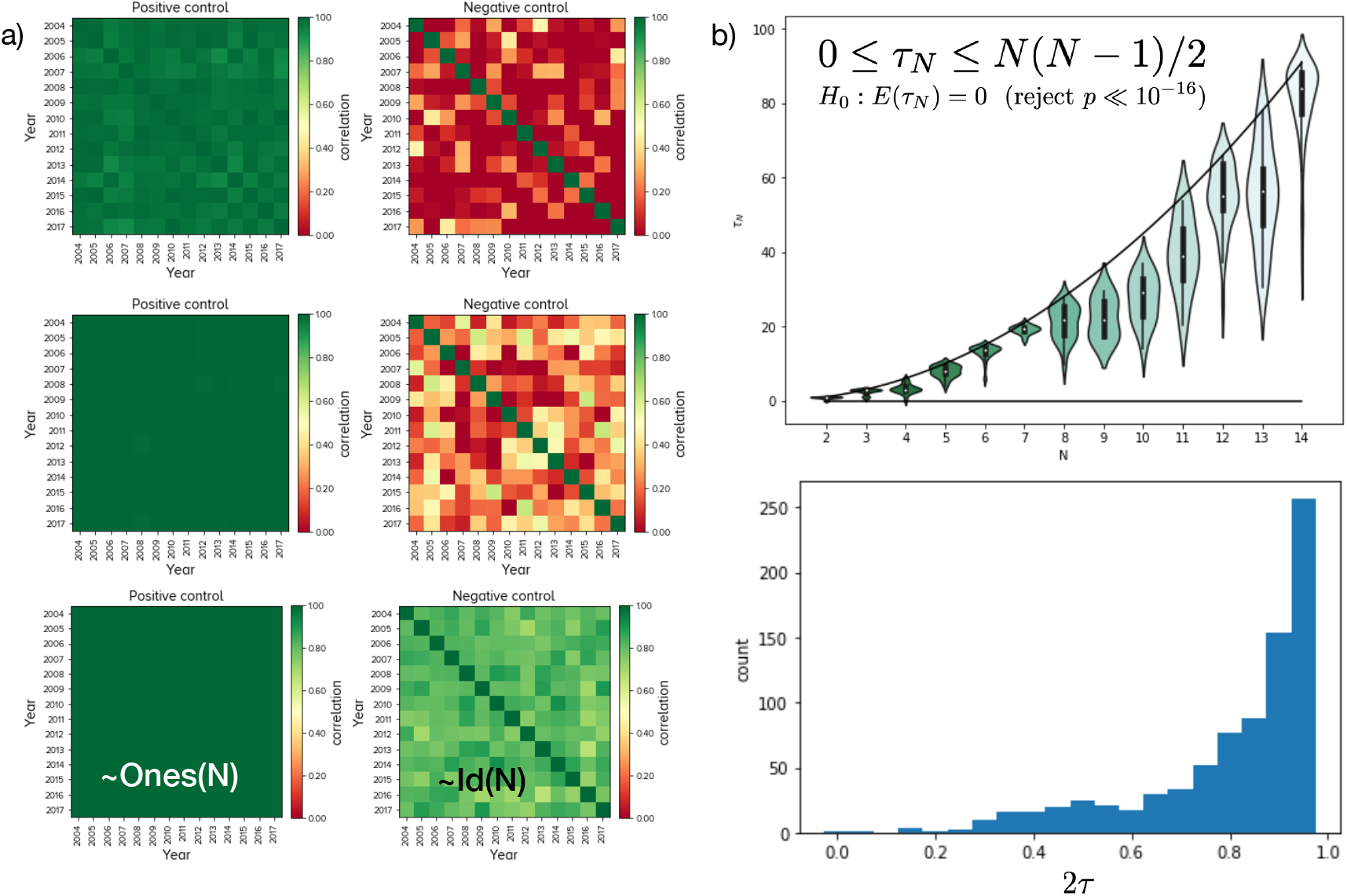
Statistics of *τ* show MIC distributions exhibit significant year-year correlations. a) Synthetically generated correlation matrices from timeseries of processes both with and without temporal correlations illustrate the near-ones matrix, Ones(*N*), and near-identity matrices, Id(*N*) that result from those different processes. b) Statistics that result when *τ* is applied to correlation matrices for Atlas PA pairs having MIC timeseries for *N* years: *τ* lies somewhere between 0 and *N*(*N* – 1)/2 whereby all densities significantly are closer to the latter than the former, indicating non-random correlations. If we assume a null model whereby Atlas is ‘noise-driven’ in the sense that MIC data for all strains and all antibiotics can be modelled as i.i.d. random variables with no between-year MIC correlations, this null can be rejected with very high probability by testing that the mean of *τ* = 0.

**Fig. S6:**
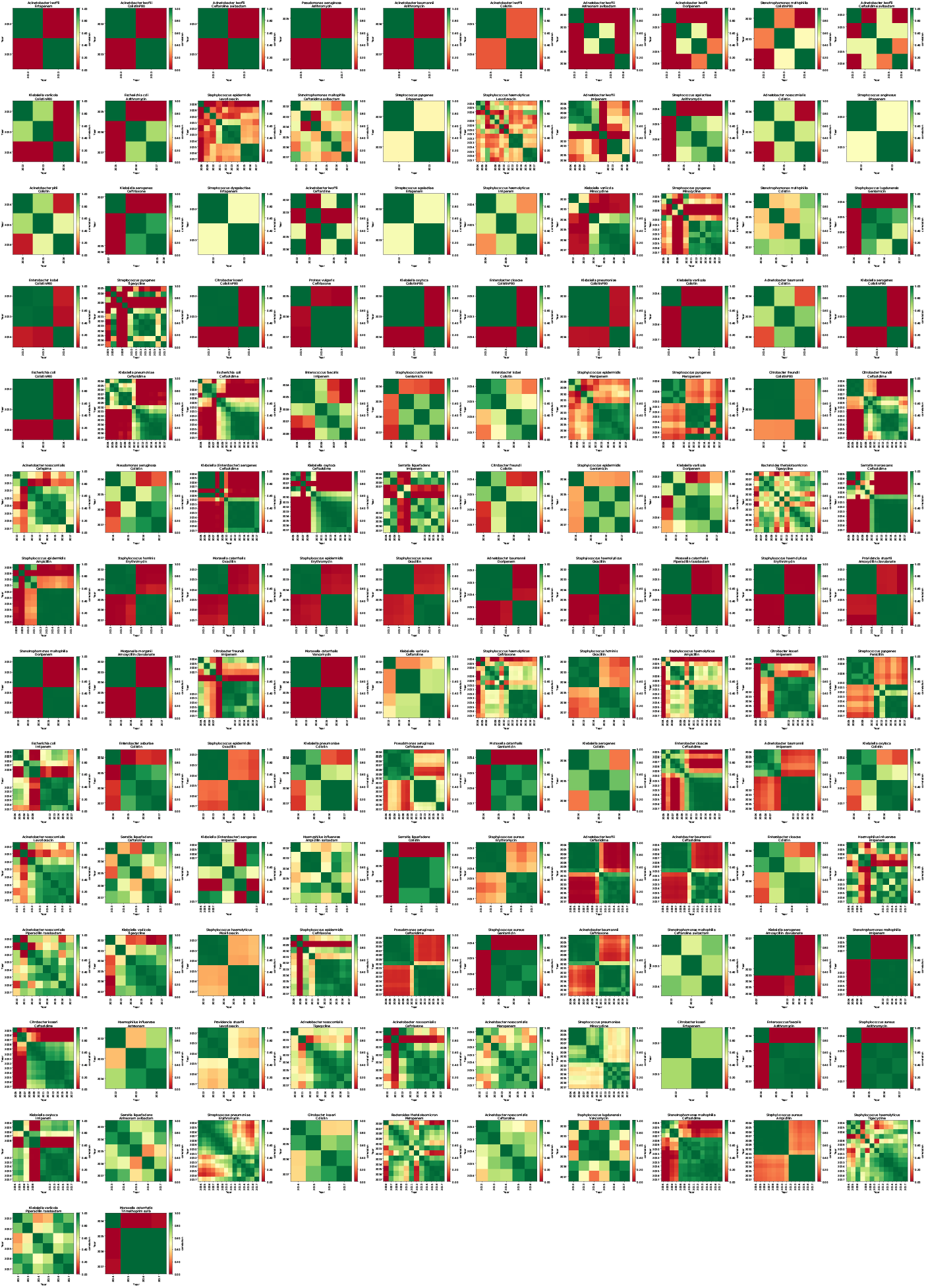
The ‘worst’ PA pair year-year correlations ranked by *τ*: these satisfy *τ* < 1/4. These MIC correlations are poorly ranked with respect to *τ* and many only have data for 2 or 3 years. Others with more data are poorly ranked because they have block structures consistent with high year-year correlations followed by sudden changes in MIC distribution.

**Fig. S7:**
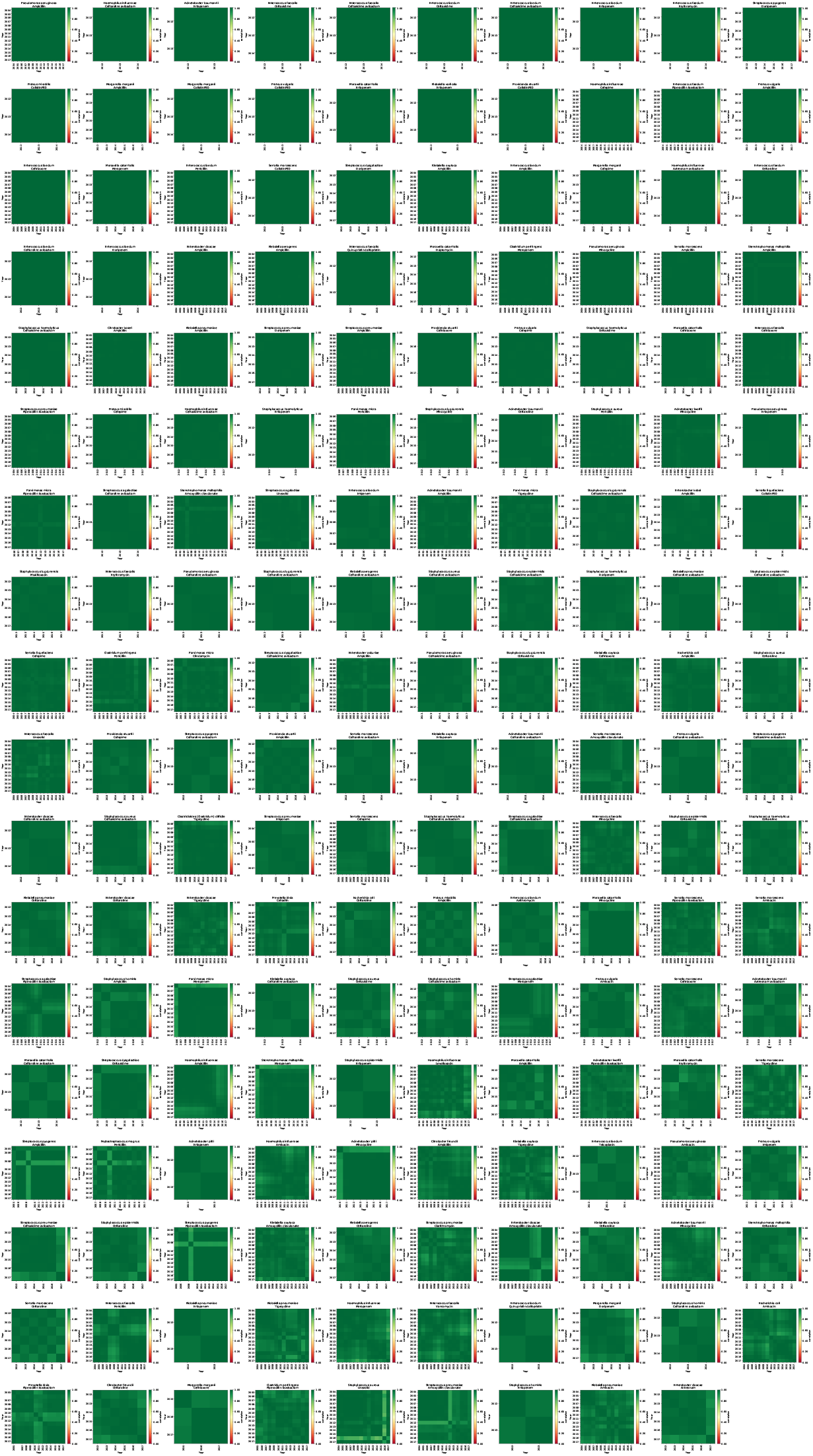
The ‘best’ PA pair year-year correlations ranked by *τ*: these satisfy *τ* > 0.485. These MIC correlations are well-ranked with respect to *τ*, they are almost monochrome, consistent with persistent year-year correlations.

**Fig. S8:**
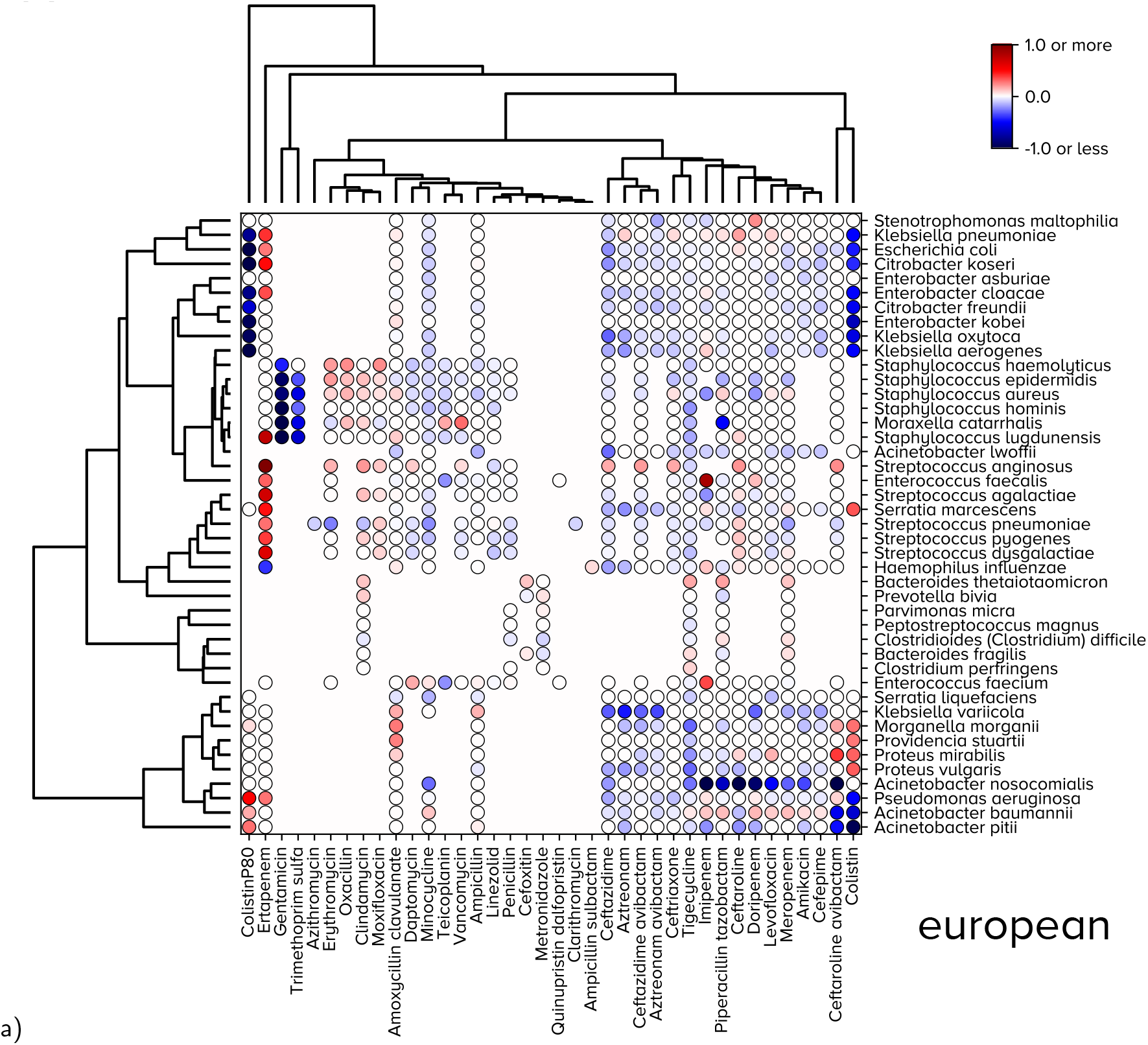
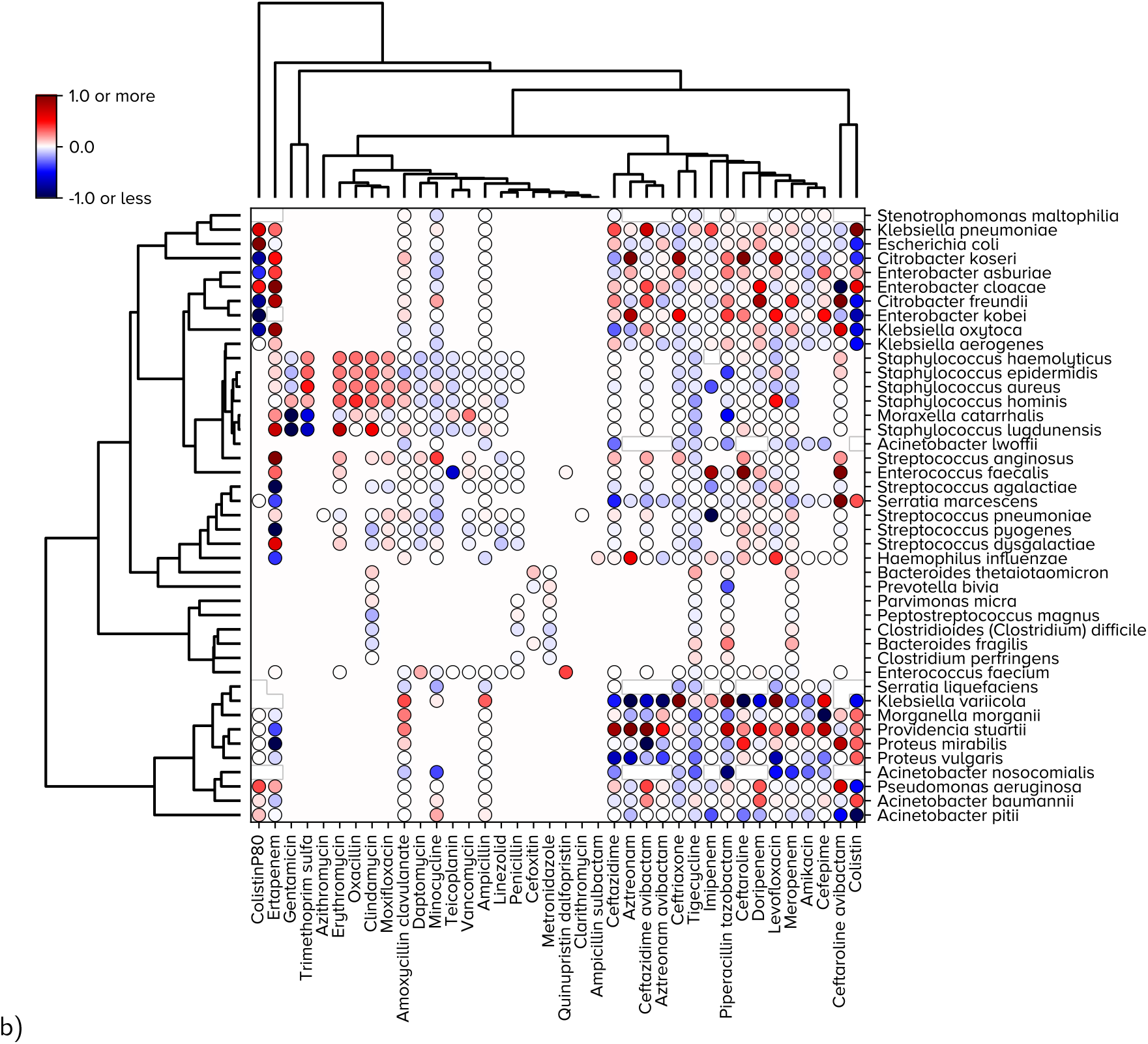
Two types of predicted change for European MIC data: mean in (a) and extreme change in (b). a) Linear regression applied to the global MIC distribution for each of the shown PA pairs indicates a motif of increases and decreases for Europe-wide mean MICs where the colour is the rate of change estimate provided by the regression’s slope. b) However, comparing a) with the analogous linear regressions conducted on the R cluster only for each PA pair (again for European data) indicates a bias towards red: this is consistent with some increases in R even when mean MICs are decreasing.

**Fig. S9:**
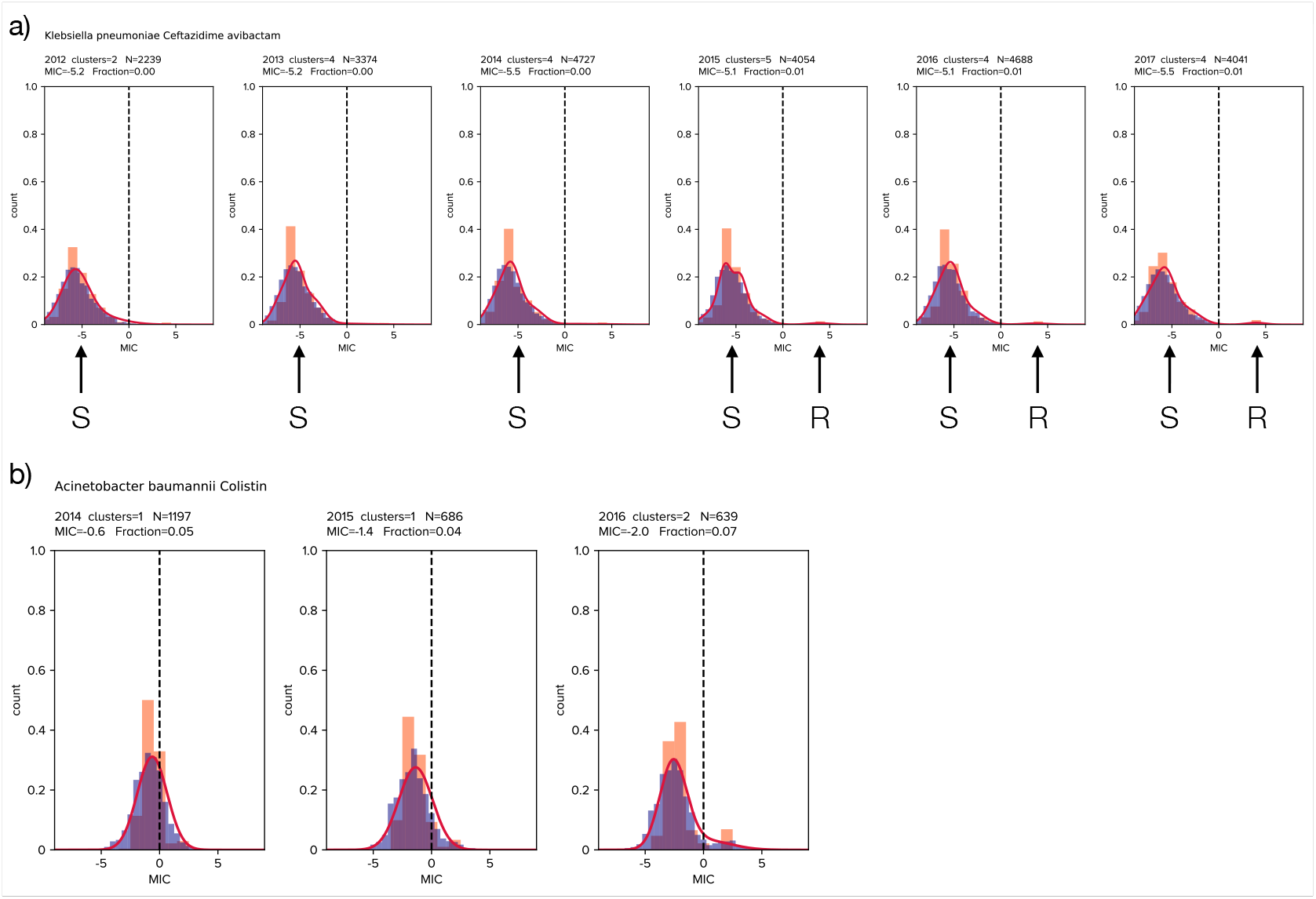
Exemplar determinations of S and R cluster pairs and their dynamics. a) A Gaussian mixture model approach does not separate the *Klebsiella pneumoniae* population into multiple clusters with different MIC phenotypes to ceftazidime avibactam prior to 2015 as its most likely model is formed from significantly overlapping Gaussians. After 2015 this changes and around 1% of the population resides in a Gaussian cluster with significantly higher mean MIC than the remainder; at this time the high-MIC cluster is labelled R and the remainder is labelled S. This resembles the clinical procedure of allocating pathogens using a binary classification, S and R, based on breakpoints whereas this cluster modelling approach determines the R-classification breakpoint as part of an algorithmically-defined procedure. Performing the clustering longitudinally identifies R clusters that are sub-clinical and yet which have increasing MICs, so they could become clinically resistant. b) This is analogous to a) for *Acinetobacter baumanii* and collistin over a 3-year period. NB: orange bars are histograms of Atlas MICs, blue histograms include modelled uncertainty and red lines are estimated kernel densities determined from Gaussian mixtures.

**Fig. S10:**
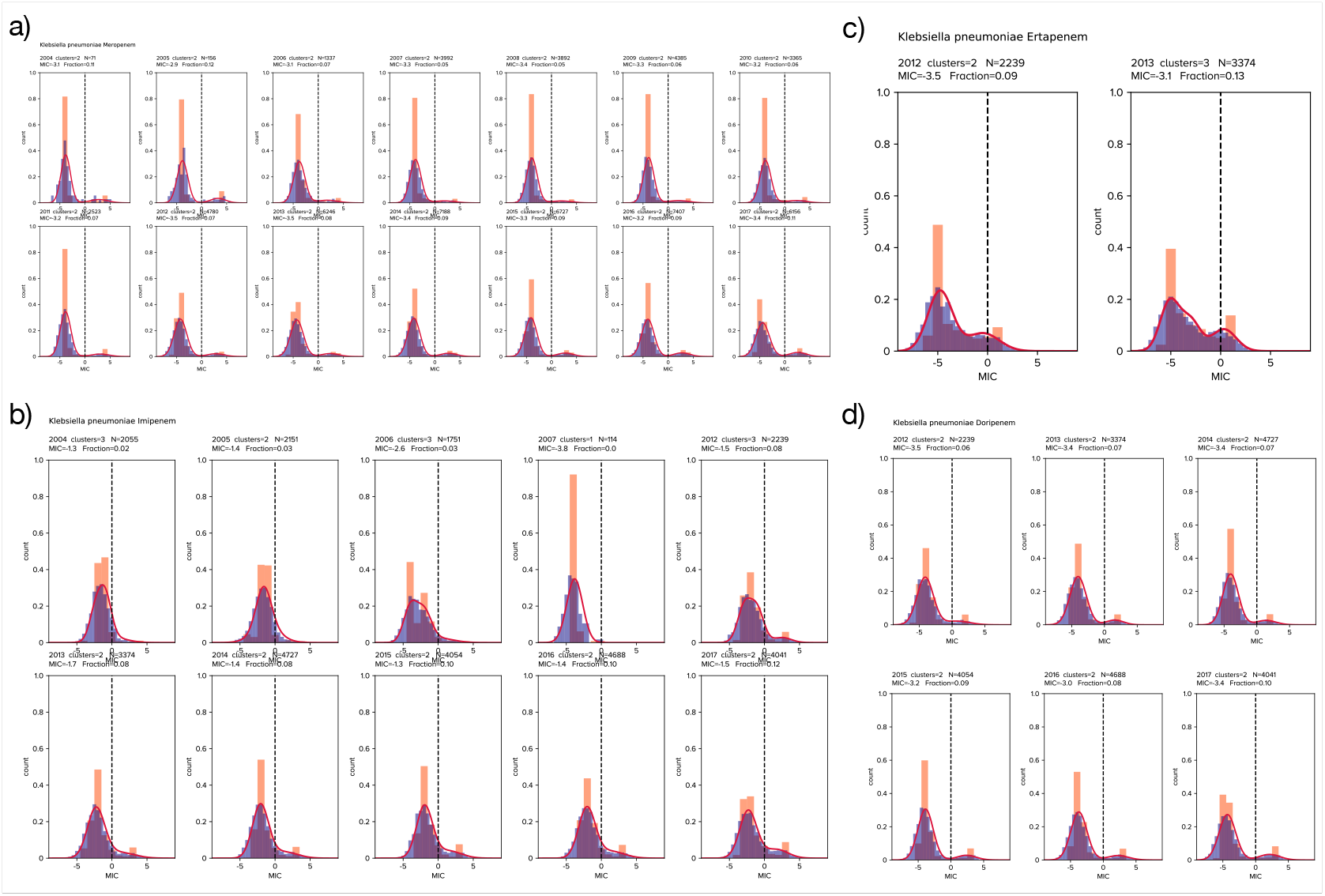
Emergent and stable bimodal SR clusters for *Klebsiella pneumoniae* and carbapenems: a) meropenem; b) imipenem - note the emergence of a clinically resistant R cluster from 2006; c) ertapenem; d) doripenem. (NB: orange bars are histograms of MICs, blue histograms include modelled uncertainty and red lines are kernel densities from Gaussian mixture models.)

**Fig. S11:**
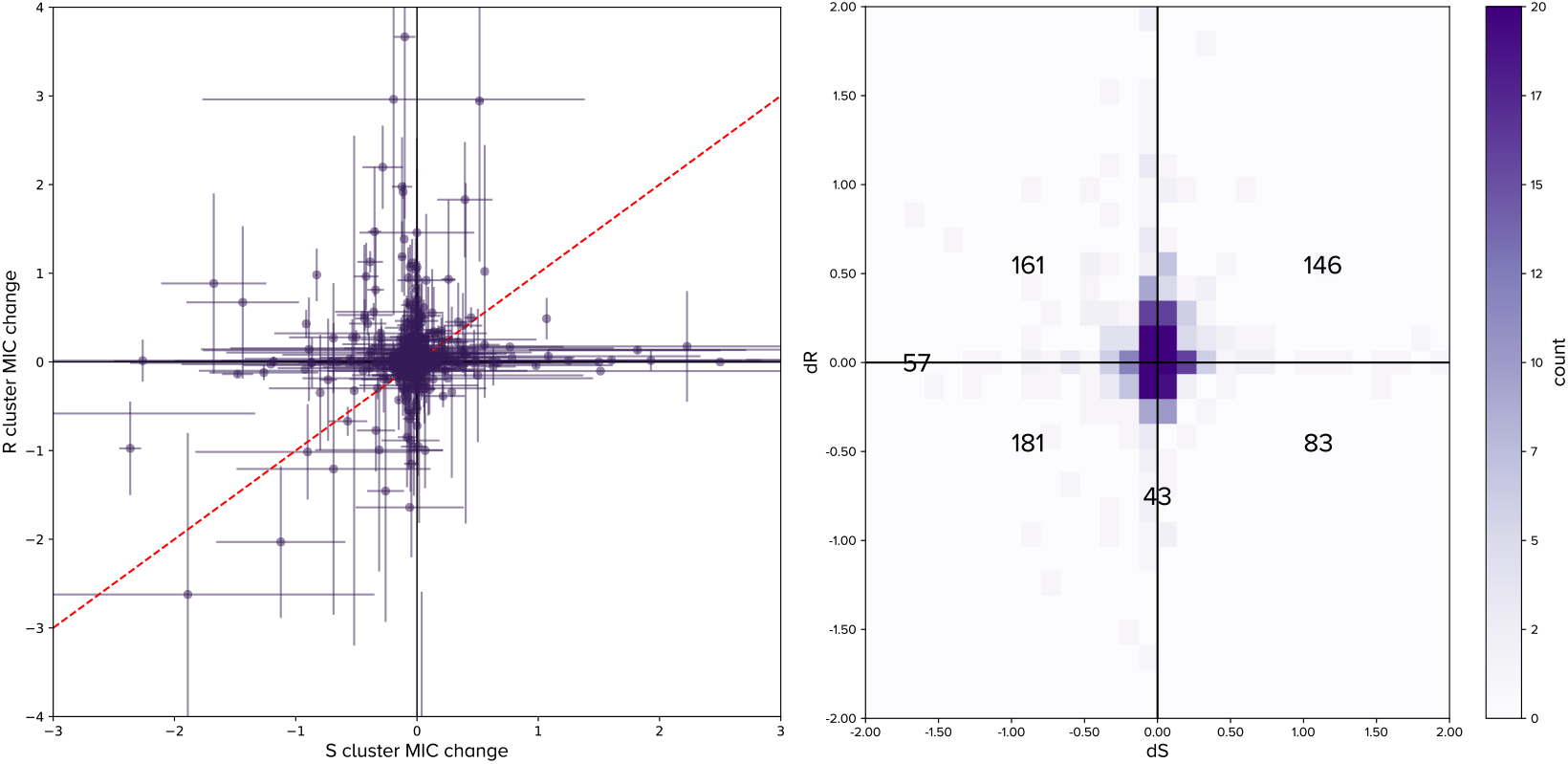
The phase plane of time derivatives of the mean MICs of S and R clusters for PA pairs. The rate of change of S and R for each PA pair on the left hand plot is shown as a dot (one standard deviation, *n* = 50) and the right hand plot counts those on the left that reside significantly in each quadrant, or else they lie on the boundary between two quadrants. We identify disruptive selection as being PA pairs with increasing MIC derivative in the R cluster but decreasing MIC derivative in the S cluster, these are points in the top-left quadrant. Purifying selection occurs when the MIC of S is increasing but that of R is decreasing, these are points in the bottom-right quadrant.

**Fig. S12:**
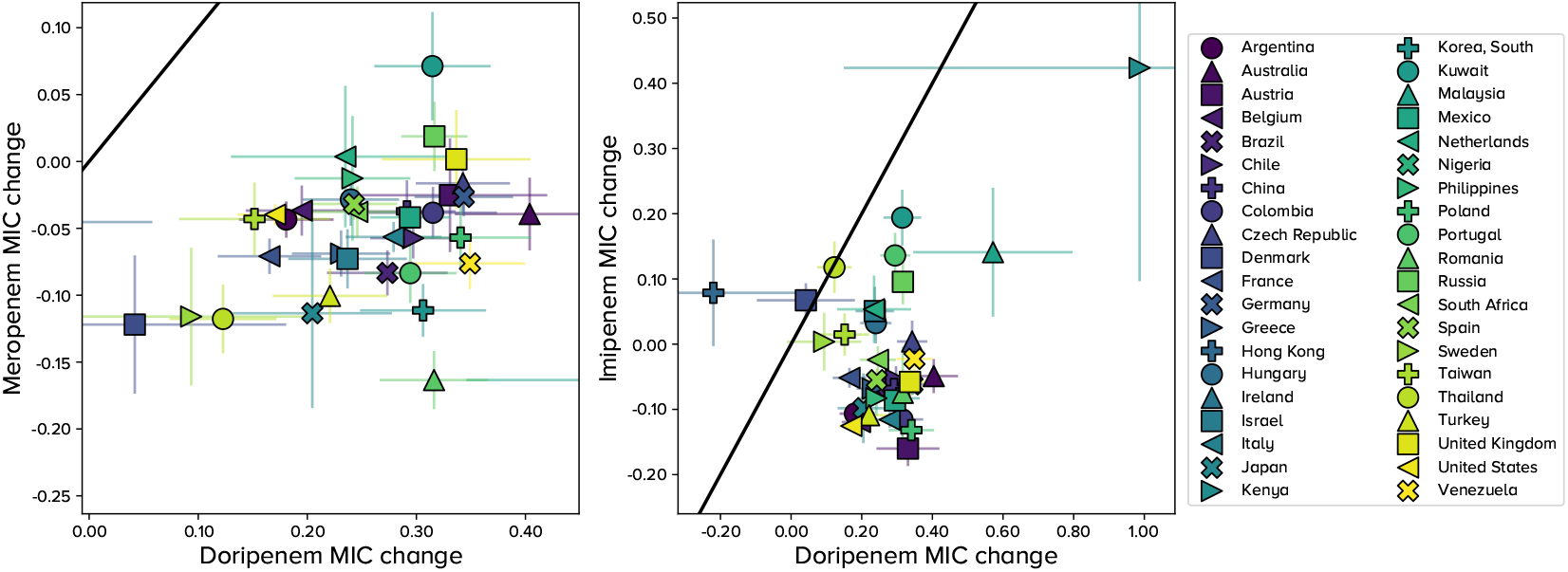
Time derivatives of the mean MIC of R for *Pseudomonas aeruginosa* and 3 carbapenems. The black line denotes ‘*y* = *x*’ in both plots, thus indicating countries for which doripenem MICs change at the same, or different, rates as meropenem and imipenem: doripenem has more rapidly increasing R MIC, independently of the country analysed. Bars represent standard error of the mean, *n* is different for each. This analysis was performed on one synthetic replicate of Atlas data (see Methods).

**Fig. S13:**
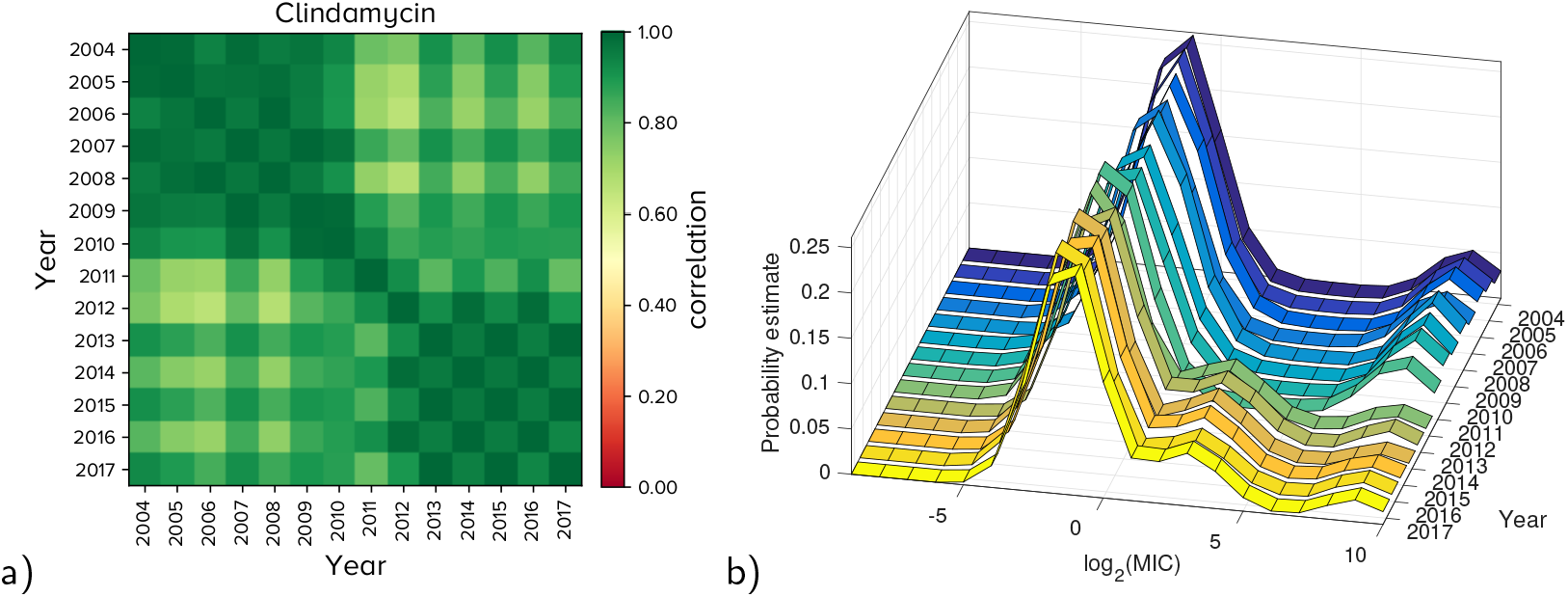
Shifting MIC dynamics of PA pair *S. pneumoniae* and clindamycin. Compare this with Figure 1C which shows a very similar structure in the analogous figure for *S. pneumoniae* and erythromycin. Note the correlelogram in a) and the temporal dynamics of the MIC distribution in b) which shows a structural shift in 2010-2011.

